# Specificity of California mouse pup vocalizations in response to olfactory cues

**DOI:** 10.1101/2021.10.12.464084

**Authors:** Kerianne M. Wilson, Victoria Wagner, Wendy Saltzman

## Abstract

In rodents, young pups communicate with their parents through harmonic calls and ultrasonic vocalizations (USVs). These forms of communication can improve chances of survival, since pups rely on their parents for thermoregulation, nutrition and protection. The extent to which pups modulate calls in response to their surroundings remains unclear. In this study we examined whether olfactory stimuli influence characteristics of pup calls, and how these calls may be affected by pup sex and litter size, in the California mouse (*Peromyscus californicus*). Pups were isolated and audio recorded during an initial, 3-minute control period, after which they were exposed for 5 minutes to bedding containing one of 4 olfactory cues: scent from their home cage, scent from the home cage of an unfamiliar family, coyote urine, or no scent. Latency to call, call rate, call duration and call characteristics (e.g. frequency and amplitude) were compared between the control period and olfactory-exposure period as well as among olfactory conditions. Pups from 2-pup litters called more quietly (lower amplitude) when exposed to odor from a predator while pups from 3-pup litters called louder (higher amplitude). Additionally, pups tended to reduce their call rates in response to odors from their home cage, consistent with contact quieting. However, pups tended to increase their rate of calling when exposed to predator urine, in contrast to the expectations of predator-induced vocal suppression. Lastly, male pups produced higher-frequency calls and more USVs than females. These results indicate that a number of pup call characteristics in this species can be influenced by acute olfactory stimuli as well as factors such as litter size and sex. The value of these pup call variations for offspring-parent communication is unclear: whether they elicit different parental responses is unknown and would be an interesting/valuable/informative avenue for future studies.

## INTRODUCTION

Vocalizations are a key component of rodent social behavior, including parent-offspring communication. Pup calling behavior in several species of rodent consists of ultrasonic vocalizations (USVs) and is vital for survival of dependent young, as it reliably elicits parental care (reviewed in Hofer et al. 2001, Shair 2018a). Pup vocalizations show consistent patterns in response to stimuli, which has allowed this behavior to become a model for studying, among other things, infant distress (Lingle et al 2012; Esposito et al. 2017) and autism spectrum disorders (Scattoni et al. 2008; Wohr and Scattoni 2013). For example, several studies report parental potentiation (an increase in a pup’s production of isolation calls after a reunion with its mother or family; reviewed in Shair 2007, 2014) and higher call rates in cold rather than thermoneutral environments (Okon 1970, 1971; Allin and Banks 1971; De Ghett 1974; Oswalt and Meier 1975). While these studies provide insight into general functions of pup vocalizations, much remains to be understood about the type of information a pup may convey when it calls. A better understanding of how and why pups may alter their vocalizations as a result of both organismal and environmental condition, such as litter size, sex and ecologically relevant olfactory stimuli, might improve the use of USVs as a model for infant communication and behavior.

Pup vocalizations are likely to be under selection to remain flexible in order to maximize their communication value (Tibbetts and Dale 2007; Kalcounis-Rueppell et al. 2018). Therefore, acute variation in a pup’s surroundings will likely inform calling decisions, with pups altering, for example, the rate, amplitude (loudness), or frequency (pitch) at which they call in response to environmental stimuli. Chemosensory stimuli are likely to be especially salient: rodent chemosensory systems develop prenatally, whereas pups’ eyes and ears do not open until several days to weeks after birth (Geyer 1979, Heyser 2004). Thus, young pups must rely on olfaction to navigate and vocally respond to their environment (e.g., Schaal 2010; Ain et al. 2013; Shair 2018).

Pup vocalization rates are thought to reflect the animal’s internal state, where increased calling indicates distress and decreased calling indicates relaxation (Schwarting and Wöhr 2012), although discrepancies in these patterns have been reported (Wiedenmayer and Barr 1998). Exposing isolated mouse or rat pups to familiar nest scents significantly decreases vocalization rates, similar to contact quieting (Oswalt and Meier 1975; Geyer 1981; Verjat et al. 2019; reviewed in Shair 2007). In contrast, exposing pups to predator odor has been found to either increase (Verjat et al. 2019) or have no effect on calling rates (Wiedenmayer and Barr 1998). Potential explanations for these contrasting findings on responses to predator odor include differences in pup age and environmental temperature: younger pups and pups in thermoneutral temperatures display a reduced response to predator odor compared to older pups or pups in cold environments (Wiedenmayer and Barr 1998).

Sex and litter size have also been found to influence pup vocalizations in rodents. Early work showed that pups call at very low rates when in contact with other pups, regardless of the presence of their mother (Hofer and Shair 1987), and in some species, pups from small litters call less than those from larger litters when isolated (prairie and pine voles [Blake 2012]; neotropical singing mice [Campbell et al 2014]). Sex differences in pup vocalizations have also been reported, but patterns are not consistent across species (e.g., Hahn et al. 1998, Vieira and Brown 2002; Wright and Brown 2004). Sex differences in pup calling are thought to reflect differences in developmental rates between the sexes, resulting from hormonal differences (Wright and Brown 2004), and may help parents discriminate among offspring based on sex (Hahn et al. 1998; Siemers et al. 2005), although olfactory cues also play a significant role in pup sex discrimination (Moore 1981; Moore and Samonte 1986). If rodent parents are able to distinguish between male and female offspring, they may differentially invest in their pups based on, for example, the pups’ reproductive value (Rutkowska et al. 2011; McGuire et al. 2016).

Because pup calls reflect distress (Schwarting and Wöhr 2012), variations in predator-induced vocalizations may reflect variations in stress experienced before and during exposure to a predator cue. For example, adult rats produce more calls during fear conditioning if they experienced stress as a pup (Yee et al 2012), and responses to fear conditioning are associated with individual differences in anxiety (Borta and Schwarting 2005). Given the potential for pup sex and litter size to influence calling behavior in pups, it may be beneficial to consider these variables concurrently with effects of olfactory stimuli on pup vocalizations.

The monogamous, biparental California mouse (*Peromyscus californicus*) presents a valuable system in which to explore natural variation in pup vocalizations because pups receive significant care from both parents (Gubernick et al. 1993; Gubernick and Teferi 2000), which stands in contrast to the majority of rodent species typically used as models for pup calling behavior and may influence patterns of offspring-parent communication. Furthermore, both pup and adult vocalizations are well described (e.g., Kalcounis-Rueppell et al. 2006; Johnson et al. 2017; Hurley and Kalcounis-Rueppell 2018; Kalcounis-Rueppell et al. 2018), allowing for more nuanced analyses of vocalizations. *P. californicus* pups produce sustained vocalization (SV) calls, which are composed of multiple stacked harmonics that reach peak frequencies in the ultrasonic range (above 20kHz; reviewed in Kalcounis-Rueppell et al. 2018). Pup vocalizations follow the typical ontogenetic pattern observed in other rodent species (e.g., De Ghett 1974; Blake 2002; Campbell et al. 2014; Kaidbey et al. 2019; Zaytseva et al 2020), with the number of isolation calls increasing from birth until pups open their eyes and become mobile (Geyer 1979) (∼14-16 days of age in California mice; Johnson et al. 2017; Zhao et al. 2019), and may contain information about pup identity (Kober et al. 2007). Because this species produces small litters (1-4 pups; mean litter size = 2; Gubernick and Alberts 1987; Cantoni and Brown 1999), naturally occurring differences in litter size may be more likely to affect parental care and offspring development than in species with larger litters.

This study assessed the effects of ecologically relevant olfactory stimuli on pup vocalizations and evaluated whether these effects were modulated by the pup’s sex and litter size. Because males of many species cannibalize pups, adult males or bedding from adult males’ cages are often used as predator cues and can result in predator-induced suppression of pup vocalizations (Hofer et al. 2001). However, rat pups reared in the presence of their sire or the sire’s odor exhibit contact quieting rather than the predator-induced response (Brunelli et al. 1998; Shair 2007). For naturally biparental species, it is unclear whether odors from adult males elicit behavioral inhibition, so cues from heterospecific predators are preferred stimuli. We used coyote urine as a predator odor in this study because adult California mice are highly sensitive to this stimulus, which elicits a large increase in plasma corticosterone concentrations (Chauke et al. 2011; Harris and Saltzman 2013). We also examined pups’ vocal responses to olfactory stimuli from an unfamiliar family, to determine whether pups are able to distinguish between the scent of their own family and a novel family. We predicted that 1) pups with fewer siblings would call less than pups with more siblings, 2) in line with previous reports for this species (Vieira and Brown 2002; Wright and Brown 2004), females would call at higher rates than males, and 3) pups would display call patterns consistent with contact quieting in response to odor cues from their family but not from an unfamiliar family, and with predator-induced suppression of pup vocalizations in response to odor cues from a coyote.

## METHODS

### Animals

We used California mice that were descended from mice purchased from the Peromyscus Genetic Stock Center (University of South Carolina, Columbia, USA) and were bred at the University of California, Riverside (UCR). Families (parents and dependent pups) were housed in 44 × 24 × 20 cm polycarbonate cages with aspen shavings for bedding and cotton for nesting material and had *ad libitum* access to food (Purina 5001 Rodent Chow) and water. The ambient temperature was maintained at approximately 23 °C, humidity was at approximately 65%, and lights were on a 14:10 h cycle (lights on at 2300 h). All procedures were approved by UCR’s Institutional Animal Care and Use Committee and were conducted in accordance with the recommendations of the *Guide for the Care and Use of Laboratory Animals*.

Mice were housed as breeding pairs with their youngest litter of pups; pups were weaned at 27-31 days of age, prior to the birth of younger siblings. We used two pups from each of 36 multiparous breeding pairs. Only one litter, containing either two or three pups, was used from each pair; as described above, California mice give birth to litters of 1-4 pups, with a modal litter size of 2-3 (Gubernick and Alberts 1987; Cantoni and Brown 1999). All recordings were conducted on postnatal day 3-5 (day of birth = postnatal day 0), with pups from the same litter tested on the same day.

### Recording Trials

Pup vocalizations were recorded between 0700 h and 0930 h over a 3-month span. Each pup was tested under one of four treatments: exposure to 1) aspen shavings (i.e., bedding) from the pup’s own family, 2) shavings from an unfamiliar conspecific family containing age-matched pups, 3) shavings doused with predator (coyote) urine, or 4) clean shavings (control). Pups were assigned to treatments randomly except that siblings did not receive the same treatment, to control for potential litter effects. Recordings were conducted in a lighted sound-reduced chamber (internal dimensions: 58 × 62 × 62 cm). An ultrasonic recording microphone (miniMIC-FG USB Microphone, Binary Acoustic Technology; Tucson, AZ, USA) was centered 8 cm above an open-top 12 × 7.5 × 5.25 cm polycarbonate cage containing a thin layer of fresh aspen shavings. Because California mouse pups at 4-6 days of age cannot regulate their own body temperature (Gubernick and Alberts 1987; Gubernick and Teferi 2000; Rosenfeld et al. 2013) and rodent pups may adjust vocalization patterns as a means to thermoregulate when isolated (Allin and Banks 1971; Geyer 1979), the cage was placed on a heating pad maintained at 35-37°C.

Olfactory stimuli were prepared in individual air-tight containers 3-5 minutes prior to being introduced to the focal pup. Conspecific stimuli (shavings from the pup’s own family or from an unrelated family with age-matched pups) were collected from bedding directly adjacent to each family’s nest, with care taken to collect only shavings without other nesting material (cotton) or feces. Shavings were in each family’s cage for at least 24 h prior to collection. Predator stimuli were prepared by spreading 1ml of coyote urine (PredatorPee®, Main Outdoor Solutions, Hermon, ME, USA) onto 1 tablespoon of shavings and shaking the container vigorously to coat all shavings. To minimize the amount of predator scent released into the testing room, predator scent was prepared outside of the vivarium using gloves, which were replaced before leaving the preparatory room. Additionally, trials using predator scent were conducted last on each day of testing to eliminate the possibility of lingering predator scent interfering with other treatment trials. Pups that were exposed to predator urine were promptly euthanized with pentobarbital (Fatal-Plus) following their recording trial to prevent other family members from being exposed to the odor and to eliminate the possibility that the pups would be neglected or attacked by their parents.

Before each trial, the subjects’ home cage was placed in a room adjacent to the room containing the sound-reduced chamber. A randomly selected pup was removed from its home cage and immediately placed in the center of the test cage on a heating pad inside the recording chamber. Calls were recorded for an initial 3-minute period using Spect’r III (Spectral Analysis, Digital Tuning, and Recording Software, Binary Acoustic Technology, Tucson, AZ, USA) at a sampling frequency of 250 kHz. After 3 minutes, the acoustic chamber was opened and 1 tbsp of stimulus shavings was placed inside the cage, directly in front of the pup. The chamber was quickly closed, and the pup was recorded for an additional 5 minutes. The pup was then removed from the acoustic chamber and its front right paw was marked with a Medline skin marker for identification; the pup was then returned to its home cage. Pups were away from their home cage for less than 10 minutes. The acoustic chamber was cleaned with Virkon disinfectant, which is used regularly in the vivarium, and the test cage was replaced by a clean cage, which was given time to heat up.

The first pup tested was reunited with its family for at least 10 minutes before the second pup was removed for testing. The trial for the second pup followed the same procedure as the first but with a different olfactory stimulus, and the back right paw of the second pup was subsequently marked for identification. After the second pup was returned to its home cage, the family was observed for at least 5 minutes to confirm normal parenting behaviors. Call characteristics (latency to first call, mean call amplitude, mean peak frequency, number of USVs, and total number of calls) did not differ between pups tested first or second (independent t-tests: *P*’s > 0.11).

To allow continued identification of individual pups, each pup’s front or back foot was re-marked every 3-4 days until we could confidently determine pups’ sexes on postnatal day 14-24 based on the presence (female) or absence (male) of nipples (Deeny et al. 2016). Sex of pups euthanized following exposure to coyote urine was determined by dissecting pups to observe the testes or uterine horns.

### Acoustic Analyses

Pup vocalizations were analyzed with Avisoft - SASLab Pro software (Avisoft Bioacoustics, Berlin, Germany). Visual inspection of sonograms revealed that all pup calls produced were sustained vocalizations. Thus, vocalization data could be averaged across all calls for a given time period. The following call characteristics were collected from each recording: latency to initiate vocalizations (call latency (s)), average call loudness (call amplitude (dB)), average maximum call pitch (peak frequency (Hz)), total number of calls, and total number of calls in the ultrasonic range (peak frequency above 20 kHz).

### Statistical Analyses

Analyses were performed in STATA 15 (StataCorp LP, College Station, TX, USA). Assumptions for linear mixed-effects models (LMMs) and t-tests were checked by evaluating quantile-quantile plots and through Shapiro-Wilkes analyses. Call latency data were square-root transformed to meet assumptions; all other data met assumptions for parametric tests and therefore were not transformed for analysis. Significance was assessed based on α = 0.05 (two-tailed).

Because acoustic recordings varied slightly in length, standardized segments of all recordings were analyzed (Fig. 1). Paired t-tests were used to assess temporal changes in call characteristics (the first 150 s of isolation versus the first 150 s following olfactory stimulus introduction), using only the pups in the control condition (i.e., exposed to clean shavings) (Fig. 1: Early vs. Late).

**Figure 1:**
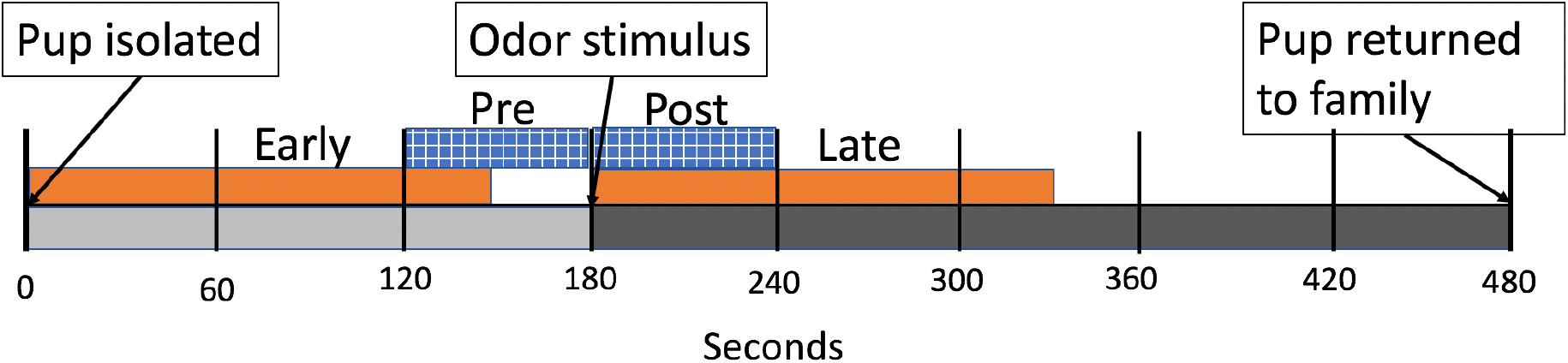
Timeline of acoustic recording procedure for each pup. Light grey shading indicates the period of isolation before the olfactory stimulus (aspen shavings that were clean, from the pup’s own family, from an unrelated family, or scented with coyote urine) was introduced, and dark gray indicates the period of isolation after the olfactory stimulus was introduced. Orange indicates the time window of each acoustic recording used to assess how pup-call characteristics change over time in isolation (Early vs. Late) and how call characteristics are influenced by pup sex and litter size (Early). Blue indicates the time windows used to assess the effects of olfactory stimuli and interactions with pup sex and litter size on pup-call characteristics (Post – Pre).

LMMs were used to assess the effects of pup sex, litter size and their interaction on characteristics of calls produced in the first 150 s of isolation (prior to introduction of olfactory stimuli), using pups in all four stimulus conditions (Fig. 1: Early). Litter identity was included in models as a random affect. Non-significant (*P* > 0.05) interactions and then non-significant main effects (pup sex and litter size) were removed from models in a reverse step-wise manner.

Lastly, LMMs were used to assess the main effects of olfactory stimulus, pup sex, litter size and the interactions between 1) olfactory stimulus and pup sex and 2) olfactory stimulus and litter size on characteristics of calls. For these analyses, we used the difference between the 60 s immediately following the introduction of the olfactory stimulus and the 60 s immediately preceding the introduction of the olfactory stimulus (Fig. 1; Post – Pre). Litter identity was included as a random effect. Again, non-significant (*P* > 0.05) effects were removed in a reverse step-wise fashion beginning with non-significant interactions and then non-significant main effects with the exception of olfactory stimulus type, since this was the main manipulation of the study.

## RESULTS

A total of 72 pups were recorded. Under lab conditions, California mouse pups are almost continuously latched onto the mother’s nipples (pers. obs.). However, 6 pups from 5 litters were not latched onto the mother when we removed them from their home cage, and these pups appeared to display different call behavior compared to latched pups. Thus, recordings from these pups were excluded from the main analyses. The final sample size was 66 pups: 28 males and 38 females, which came from 18 2-pup litters and 17 3-pup litters. Compared to latched pups, unlatched pups tended to produce fewer calls during the Early (X ± SE: unlatched – 127.0 ± 61.7; latched – 224 ± 11.8; t-test: t = −1.95, *P* = 0.055) and Late recordings (unlatched – 105.5 ± 58.8; latched – 193.5 ± 10.6; t = −1.95, *P* = 0.056). Unlatched pups also tended to produce quieter calls than latched pups during both the Early (0.23 ± 0.07 vs. 0.41 ± 0.02 dB, respectively; t = −1.89, *P* = 0.063) and Late recordings (0.17 ± 0.04 vs. 0.40 ± 0.02 dB, respectively; t = −2.26, *P* = 0.027).

### Changes in Call Characteristics Over Time in Isolation

Overall, pups tended to call at high rates, usually exceeding about 70 calls per minute throughout the period of isolation. Pups produced more calls (*P* = 0.008) and more USVs (calls with peak frequencies above 20kHz; *P* = 0.029) during early isolation compared to late isolation (Table 1). After initial placement in the chamber, control pups showed a latency of 5.5 ± 1.3 s (mean ± SE) before their first call. In contrast, they showed a much shorter latency of about 1.4 ± 0.4 s after clean shavings were placed in front of them (*P* = 0.01). Neither amplitude (dB) nor peak frequency (Hz) of calls differed between the first 150 s of isolation and the 150 s following the introduction of clean shavings (Table 1).

**Table 1.** Characteristics of calls recorded from control pups in the first 150 s after isolation and the first 150 s of exposure to clean shavings (Fig. 1: Early vs. Late). Paired t-tests, df = 16, N = 17. *P*’s ≤ 0.05 are in bold. ^a^Square-root transformed (mean and SE shown are not transformed). USVs – ultrasonic vocalizations.

| Call characteristic | Early (X±SE) | Late (X±SE) | Cohen's $d$ | $t$ | $P$ |
| --- | --- | --- | --- | --- | --- |
| Latency to first call<sup>a</sup> (s) | 5.52 ± 1.30 | 1.40 ± 0.38 | 0.897 | 3.70 | **0.002** |
| Amplitude (dB) | 0.384 ± 0.046 | 0.369 ± 0.052 | 0.159 | 0.66 | 0.522 |
| Peak frequency (kHz) | 19,706 ± 558 | 19,771 ± 614 | 0.046 | -0.19 | 0.851 |
| Number of calls | 224 ± 21 | 189 ± 20 | 0.738 | 3.04 | **0.008** |
| Number of USVs | 115 ± 16 | 94 ± 17 | 0.581 | 2.40 | **0.029** |

### Effects of Pup Sex and Litter Size on Call Characteristics

Calls during the early isolation period (first 150 s) were analyzed across all trials to assess the effects of pup sex and litter size on pup-call characteristics. Females produced calls with lower average peak frequencies compared to males (*P* = 0.05; Fig. 2A). Although pup sex did not affect the total number of calls produced in the first 150 s of isolation, females produced fewer calls with peak frequencies in the ultrasonic range compared to males (*P* = 0.034; Fig. 2B). Pup sex, litter size and their interaction did not significantly influence any other call characteristics considered here (Table 2).

**Figure 2.**
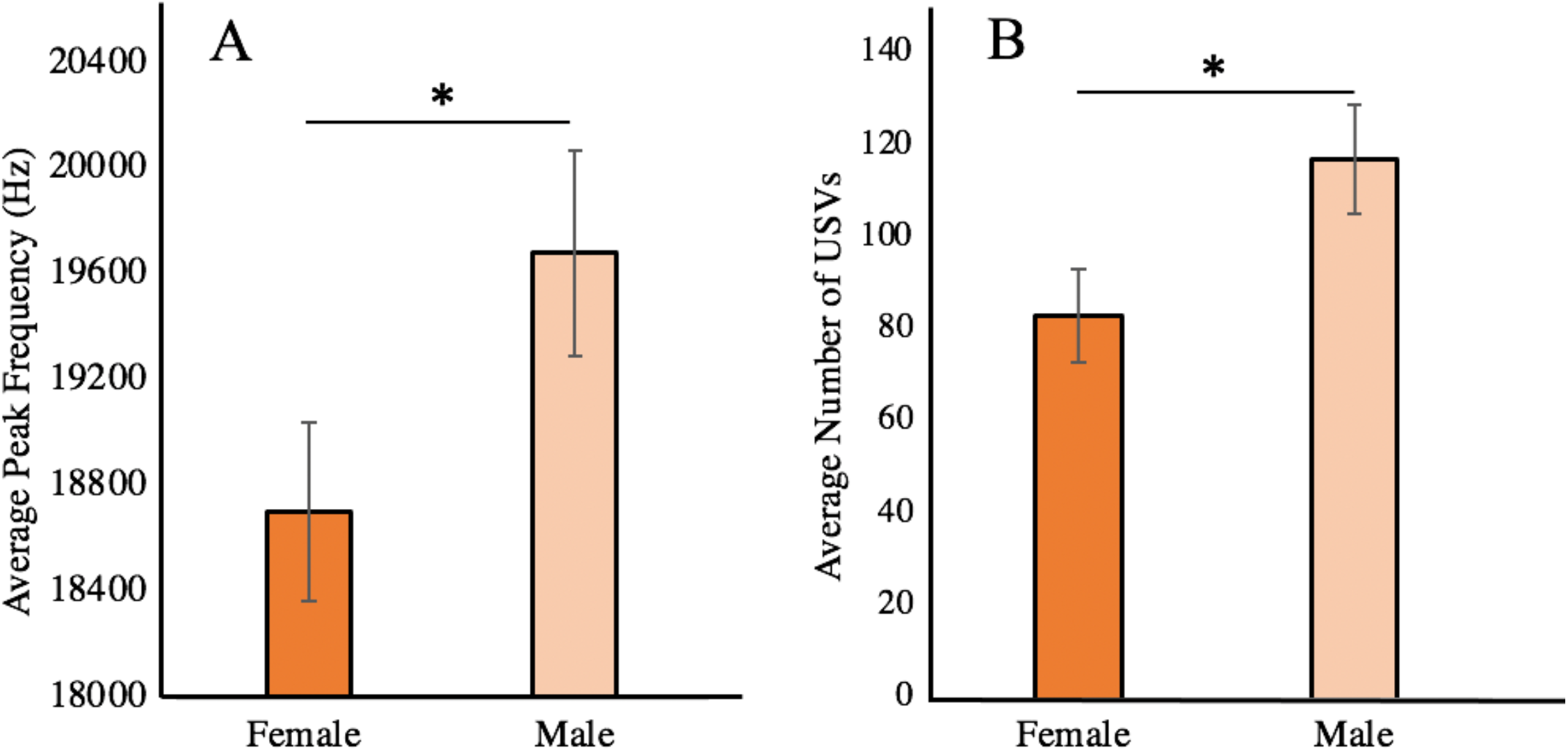
Effects of pup sex on average call peak frequency and number of USVs (calls above 20kHz) (non-transformed mean ± SE) produced during the first 150 s in isolation, prior to introduction of olfactory stimuli. Females: N = 38; Males: N = 28. Data correspond to analyses in Table 2. \**P*’s ≤ .05.

**Table 2.** Effects of pup sex and litter size on characteristics of pup calls during the first 150 s of isolation, before olfactory cues were introduced (Fig. 1: Early). Linear mixed-effect model, N = 66 pups from 35 litters. *P*’s ≤ 0.05 are in bold. ^a^Square-root-transformed. ^b^Litter identity contributed significantly to the model.

| Call characteristic | Coefficient ± SE | 95% CI | $\chi^2$ | *z* | *P* |
| --- | --- | --- | --- | --- | --- |
| Latency (s)<sup>a</sup> |  |  | 0.88 |  | 0.348 |
| Sex | -0.201 ± 0.215 | -0.622 – 0.219 | | -0.94 | |
| Amplitude<sup>b</sup> (dB) | 0.071 ± 0.052 | -0.030 – 0.173 | 1.90 | | 0.168 |
| Litter size |  |  |  | 1.38 |  |
| Peak frequency (Hz) | -977.3 ± 498.0 | -1953.3 – 1.28 | 3.85 | | **0.049** |
| Sex |  |  |  | 1.96 |  |
| Number of calls<sup>b</sup> | 22.27 ± 20.84 | -18.58 – 63.12 | 1.14 | | 0.285 |
| Sex |  |  |  | 1.07 |  |
| Number of USVs | -33.09 ± 15.59 | -63.64 - -254 | 4.51 | -2.12 | **0.034** |
| Sex |  |  |  |  |  |

### Effects of Olfactory Stimuli on Call Characteristics

To assess the effects of ecologically relevant olfactory stimuli on calls, we compared changes in call characteristics from the 60 s immediately before to the 60 s immediately after introduction of olfactory stimuli (Fig. 1; Post – Pre) among pups in the four stimulus conditions. The most pronounced effects were on average amplitude of pup calls, which was affected by stimulus type and the interaction between stimulus type and litter size (LMM model *P* = 0.002; Table 3). Additionally, the random factor of litter identity contributed significantly to the model. Among pups from 3-pup litters, those exposed to bedding with predator urine showed an increase in their call amplitude while those exposed to all other stimuli showed a reduction in call amplitude (Fig. 3). Among pups from 2-pup litters, those exposed to bedding with predator urine showed a larger reduction in call amplitude compared to pups exposed to clean shavings. Pups from 2- and 3-pup litters differed significantly in their responses to bedding from a conspecific family and to predator odor. In response to bedding from a conspecific family, pups from 2-pup litters showed an increase in call amplitude while pups from 3-pup litters decreased their call amplitudes. In response to shavings mixed with coyote urine, pups from 3-pup litters showed an increase in call amplitude while pups from 2-pup litters decreased their call amplitudes.

**Table 3:** Effects of olfactory stimulus and its interactions with pup sex and litter size on characteristics of pup calls, quantified as the difference between the calls produced following introduction of olfactory stimuli and immediately before the introduction of olfactory stimuli (Fig. 1: Post – Pre). Linear mixed-effect models, N = 66 pups from 35 litters. *P*’s ≤ 0.05 are in bold. ^a^Litter identity contributed significantly to the model.

| Call characteristics | Olfactory stimulus (X+SE) | | | | Coefficient ±<br>SE | 95% CI | $\chi^2$ | z | *P* |
| --- | --- | --- | --- | --- | --- | --- | --- | --- | --- |
|  | Own<br>family | Other<br>family | Predator<br>urine | No<br>scent |  |  |  |  |  |
| Change in amplitude (dB) |  |  |  |  |  |  | 22.57 | -1.02 | **0.002** |
| Litter size | | | | | -0.405 ± 0.040 | -0.118 – 0.037 | | 1.88 | 0.308 |
| Olfactory stimulus | -0.031 + 0.02 | 0.009 + 0.02 | -0.065 + 0.02 | -0.023 + 0.02 | -0.099 ± 0.036 | -0.170 - -0.027 | | 2.78 | **0.030** |
| Olfactory stimulus * litter size | | | | | 0.113 ± 0.053 | 0.009 – 0.216 | | | **0.005** |
| Change in peak frequency (Hz)<sup>a</sup> |  |  |  |  |  |  | 1.38 | 0.56 | 0.711 |
| Olfactory stimulus | -56.2 + 253.8 | 231.4 + 204.2 | -33.2+209 | 78.1 + 214.7 | -111.3 ± 276.1 | -652.5 – 429.9 | | | |
| Change in number of calls |  |  |  |  |  |  | 6.75 |  | 0.081 |
| Olfactory stimulus | -10.67 + 7.79 | -4.95 + 6.19 | 12.27 + 6.36 | -5.41 + 6.55 | 17.69 ± 9.13 | -0.204 – 35.58 | | 1.40 | |
| Change in number of USV calls |  |  |  |  |  |  | 2.97 |  | 0.396 |
| Olfactory stimulus | -0.83 + 7.37 | -2.11 + 5.86 | 11.22 + 6.02 | 1.18 + 6.19 | 10.05 ± 8.63 | -6.87 – 26.96 | | 0.26 | |

**Figure 3.**
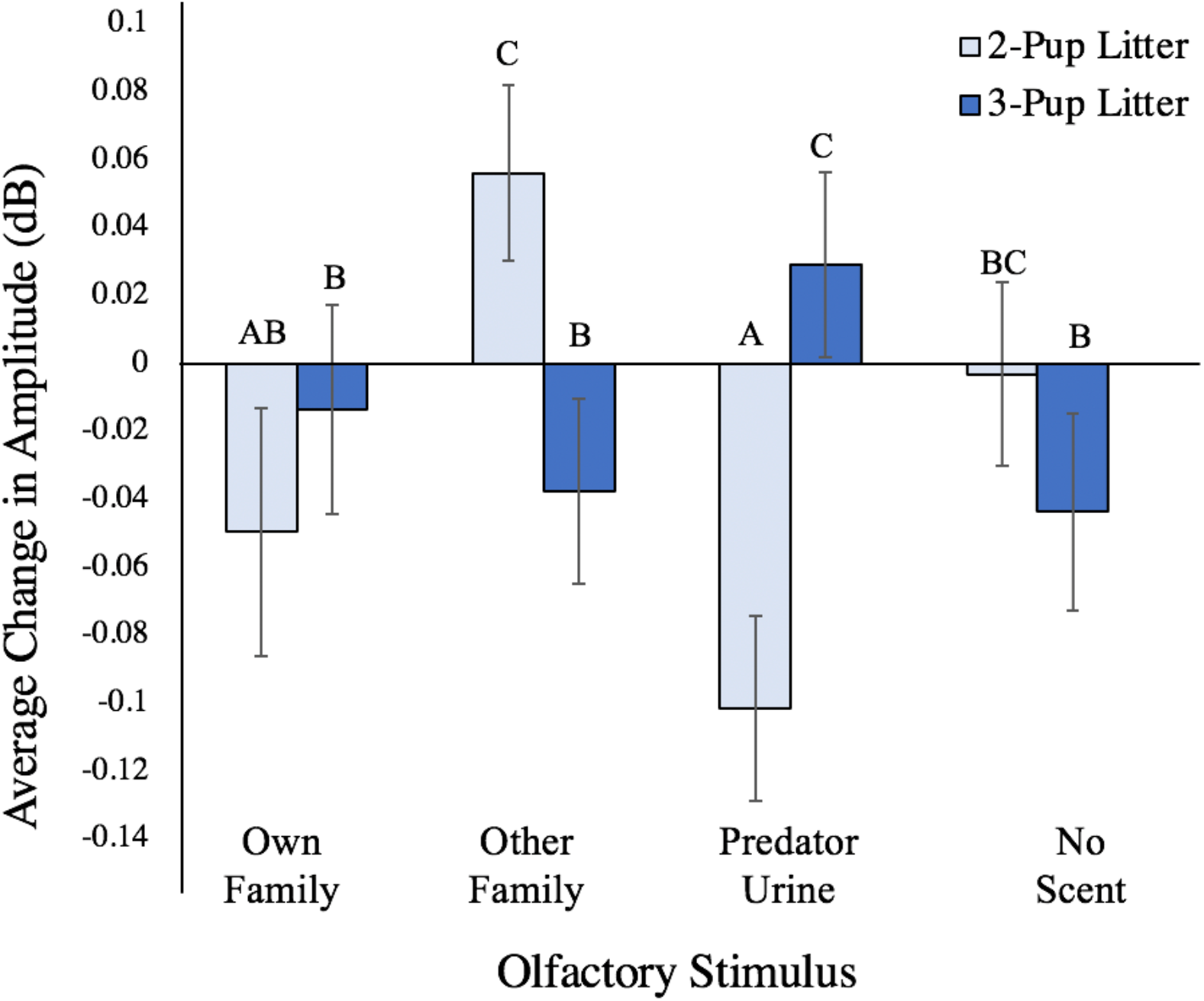
Effect of the interaction between olfactory stimulus and litter size on the change in average amplitude of calls (mean ± SE) after the olfactory stimulus was introduced. Data correspond to analyses in Table 3. Pups from 2-pup litters: N = 18; pups from 3-pup litters: N = 17. Different letters indicate significant differences among groups based on post hoc analyses. (*P*’s ≤ .05)

The change in the number of calls a pup produced showed a non-significant tendency to vary with olfactory stimuli (P = 0.081; Table 3): pups tended to show the greatest decrease in call production when exposed to bedding from their home cage and the greatest increase in call production when exposed to bedding mixed with predator urine. None of the other call characteristics that we analyzed were significantly influenced by a main effect of olfactory stimulus, an interaction between stimulus and pup sex, or an interaction between stimulus and pup age (Table 3).

## DISCUSSION

Vocalizations are the predominant means by which dependent rodent offspring communicate with their parents, and are important in eliciting parental care (Noirot 1972; Smotherman et al. 1974; Ehret and Haack 1984; Wright and Brown 2004; Portfors 2007). Here we show that this form of communication is dynamic and that variations in pup call characteristics might convey nuanced information to parents in the California mouse.

All characteristics of pup vocalizations considered here varied in response to at least one of the factors considered, including pup sex, litter size, and olfactory environment. Most notably, we found that call amplitude was jointly affected by litter size and olfactory stimulus, suggesting that a pup’s family environment informs the way in which it responds to olfactory stimuli. The trend towards an increase in calling rate associated with exposure to predator urine might indicate that pups alter their calling behavior in order to convey more nuanced information to their parents. Interestingly, call frequency was not altered by a pup’s olfactory environment, as would be predicted, but was found to differ based on pup sex.

### Temporal Effects

We assessed effects of time from the onset of isolation to gain a better understanding of the temporal changes in pup vocalizations. The initial latency until pups began to call at the start of isolation was greater than the latency to resume calling after the chamber was opened and unscented shavings were placed in the cage. In fact, we found that it was rare for pups to stop calling for more than a couple of seconds immediately following introduction of the shavings.

Surprisingly, the effects of isolation time on pup vocal characteristics are not often reported; however, the finding that pups produced fewer calls and, specifically, fewer USVs as their time in isolation progressed agrees with a previous study in prairie voles, *Microtus ochrogaster* (Lea 2006), and could reflect a response to handling stress experienced immediately before the recording began. The negative relationship between pup calling rates and time in isolation may also be related to the physiological changes that result from this behavior. Calling is energetically expensive, and rodent pups’ body temperature increases when they vocalize (Okon 1970, 1971; Allin and Banks 1971; Ghett 1974; Oswalt and Meier 1975; Blumberg and Alberts 1990; Kraebel et al. 2002). There has been some debate about whether pups call for the purpose of thermoregulation (e.g., Blumberg and Alberts 1990; Blumberg and Sokoloff 2001; Hofer 1996; Shair 2007); however, this possible function of vocalizing is unlikely to explain our results, since we maintained pups in a thermoneutral environment by placing the test cage on a heating pad.

### Sex Effects

In contrast to previous reports in this species, in which female pups were found to call more than males (Vieira and Brown 2002; Wright and Brown 2004), we found that male pups called at a higher peak frequency and, correspondingly, produced more USVs than females. The disparity between our findings and those of the earlier studies might have resulted from several methodological differences. For example, Brown and colleagues (Vieira and Brown 2002; Wright and Brown 2004) isolated pups on a heating pad before recording their vocalizations in a cold testing environment, and found sex differences at ages slightly earlier than the current study (3-4 vs. 4-6 days of age). Furthermore, the disparity in findings on sex differences in California mice is not surprising given that contradictory results have been found within other rodents as well as across rodent species. In rats, for example, female pups have been found to call both more (Brunelli and Hofer 1996) and less than males (Blumberg and Stolba 1996). Additionally, Hahn et al. (1997) found no sex difference in call characteristics in albino mice, but Hahn et al. (1998) found that females emitted fewer USVs between 4 and 5 days of age compared to males. Nonetheless, the fact that USV production differs between the sexes in several rodent species suggests that sex is an important factor and should be explored further to better understand its significance in this context.

Possible sex differences in frequency of pup calls have not been explored previously in California mice. However, a study using C57BL/6JOlaHsd and C57BL/6NCrl sub-strains of house mice found that female pups display higher frequency modulation (Wohr et al. 2008). Since pup vocalizations are influenced by parental care in house mice (Wohr and Schwarting 2008), sex differences in call characteristics may reflect differences in parental care. Rodent parents differentially invest in offspring based on sex (More and Morelli 1979; Clark and Galef 1989; Clark et al. 1990; Deviterne and Desor 1990), and the cost to parents of raising a pup differs based on the sex of the pup (Clark and Galef 1989; Clark et al. 1990). Thus, sex differences in calling frequency may provide a mechanism through which parents decipher pup sex (Kober et al. 2007) and differentially allocate care. Conversely, innate sex differences in calling could, in turn, influence parental care. The relationship between sex-dependent vocalizations and parental care has not, to our knowledge been explored. This presents a valuable area of study, which could be explored further through experiments that manipulate pup call characteristics and assess parental care.

### Litter and Olfactory Effects

This is the first study, to our knowledge, that explores modulation of pup calls in the California mouse in response to olfactory stimuli. We find support for contact quieting behavior in Californa mice, but not for predator-induced call suppression. Pups tended to reduce the number of calls they made when exposed to bedding from their home cage, which is consistent with contact quieting and has been well established in murid rodents (Shair 2007). This also agrees with previous observations of California mouse pups, which have been found to reduce calling rates when reunited with their family after a period of isolation (Johnson et al. 2017). Here, we show that home-cage bedding alone is enough to elicit this same response. Although we did not explore parental potentiation (an increase in isolated pup call rate and amplitude following reunion with parents or littermates, reviewed by Shair 2007, 2014), it is noteworthy that Johnson et al. (2017) did not find support for potentiation in this species, but contact quieting was observed.

Interestingly, our finding that pups tend to increase their calling rate in response to coyote urine counters expectations based on studies in rats, which tend to display predator-induced suppression of pup vocalizations (reviewed in Hofer 1996; Shair 2018). The disparity between pup vocal responses to predator stimuli across species may relate to the conditions under which stimuli were presented. For example, Wiedenmayer and Barr (1998) found that young rat pups fail to display predator-induced freezing behaviors when exposed to predator stimuli in thermoneutral environments. California mouse pups in our study were placed on a heating pad, which might explain why we similarly failed to observe a reduction in call rate and instead observed an increase. Finally, although we cannot rule out the possibility that the failure of young pups to reduce their calling rates may relate to an underdeveloped predator response, as suggested by Wiedenmayer and Barr (1998), the finding that pups altered their calling behavior in response to predator odor suggests that they were at least able to recognize it as a salient stimulus.

We found that litter size was an important factor influencing how California mouse pups responded to olfactory stimuli. Pups from 2-pup litters showed a greater reduction in call amplitude when exposed to predator urine than when exposed to clean bedding, while pups from 3-pup litters showed the opposite pattern. Pups from larger litters receive lower levels of parental care relative to smaller litters in meadow voles (*Microtus pennsylvanicus*) (McGuire and Bemis 2007) and red squirrels (*Tamiasciurus hudsonicus*) (Studd et al. 2016), and lower levels of care are associated with higher levels of stress in rats, as evidenced by epigenetic changes to a glucocorticoid receptor gene promoter in the brain (Weaver et al. 2004, 2006; Meaney and Szyf 2005). Since increases in call amplitude are often associated with higher levels of distress (Lingle et al. 2012), it is possible that pups from 2- and 3-pup litters differ in their affective, and therefore vocal, responses to olfactory stimuli and that this difference might result from differences in parental care received. It is important to note, however, that effects of litter size on parental care and stress responsiveness in pups have not been examined in California mice. Exploring the mechanisms by which litter size alters pups’ vocal responses to olfactory stimuli would be an interesting avenue for future research and could provide new insights into the relationship between parental care and offspring-parent communication.

## ACKNOWLEDGEMENTS

We thank Dr. Khaleel Razak and Dr. Nancy Burley for the equipment used to record and analyze pup calls, April Arquilla for technical help, Catherine Nguyen, April Arquilla, Melina Acosta and the UCR vivarium staff for assistance with colony maintenance. This research was funded by NSF grant NSF DBI-1907268.

